# Peptide Driven Identification of TCRs (PDI-TCR) reveals dynamics and phenotypes of CD4 T cells in tuberculosis

**DOI:** 10.1101/2025.05.06.652535

**Authors:** Rashmi Tippalagama, Raphael Trevizani, Leila Y Chihab, Ashu Chawla, Kai Fung, Jason Greenbaum, Kendall Kearns, Aruna D. De Silva, Wathsala Gunasinghe, Judy Perera, Hansani Gunasekera, Darsha D Senevirathne, Thomas J Scriba, Alessandro Sette, Cecilia S Lindestam Arlehamn, Julie G Burel, Bjoern Peters

## Abstract

Assigning antigen specificity to T cell receptor (TCR) sequences is challenging due to the TCR repertoire’s diversity and the complexity of TCR:antigen recognition. We developed the **<u>P</u>**eptide-**<u>D</u>**riven **<u>I</u>**dentification of **<u>TCR</u>**s (PDI-TCR) assay that combines *in vitro* expansion of cells with peptide pools, bulk TCR sequencing, and statistical analysis to identify antigen-specific TCRs from human blood. A key feature of PDI-TCR is the ability to distinguish true antigen-specific TCR clonotypes from TCRs associated with unspecific bystander activation by comparing responses to non-overlapping peptide pools. We applied PDI-TCR to Tuberculosis (TB) patients, sampling blood at diagnosis and throughout treatment, and *Mycobacterium tuberculosis* (Mtb)-sensitized healthy individuals (IGRA+). We identified hundreds of Mtb-specific TCRs, as well as unspecific TCRs, and characterized their phenotype in each cohort by single-cell RNA sequencing *ex vivo*. Mtb-specific T cells were highly diverse, with short-lived effector phenotypes only present in TB at diagnosis, while memory phenotypes were maintained through treatment. In contrast, unspecific expanded T cells were more clonally restricted, had a cytotoxic phenotype, and were maintained throughout treatment. This showcases PDI-TCR as a powerful tool for identifying antigen-specific TCRs, which enables direct *ex vivo* identification and monitoring of antigen-specific T cells.

## Introduction

Human T cells express highly variable receptors (TCRs) that recognize self and non-self antigens presented to them by Human Leukocyte Antigen (HLA) molecules on the surface of target cells. Conventional T cells express TCRs composed of an alpha (α) and beta (β) chain encoded by TRA and TRB genes that are created through a stochastic process of V(D)J gene recombination leading to a high diversity of receptor sequences that carry the ability to recognize a vast repertoire of peptide ligands^1,2^. The CDR3 segment, located within the TCR β chain, is the most variable region that directly interacts with the peptide it recognizes^1,2^. These characteristics make the CDR3 segment a key determinant for antigen recognition and, thus, the primary target in TCR sequencing^3^. Upon peptide-TCR binding, T cells can undergo expansion, resulting in a T cell population with an identical TCR, often referred to as a T cell clonotype. The ability to identify the antigen specificity of a given TCR (and the corresponding clonotypic T cell population) is important to characterize antigen-specific T cell responses during immune perturbations, such as infection, cancer or allergy^4,5,6,7^. Moreover, the ability to monitor T cell clonotypes over time and to identify the exact epitopes they recognize provides insight into the kinetics of TCR expression during disease pathology and/or protection^8,9^.

Several experimental approaches have been established to identify antigen-specific TCRs. The current gold standard is the use of fluorochrome conjugated HLA-peptide complexes (tetramers, dextramers, or spheromers) to isolate antigen-specific T cells directly *ex vivo*^10,11^, followed by single-cell TCR sequencing. This technique has been used to profile paired TCR sequences to known antigens of type 1 diabetes (T1D) and of multiple pathogens in CD8 T cells^12^. While powerful, HLA-peptide-complex based technologies are limited to known epitope/HLA combinations and can only be applied to individuals who express the corresponding HLA alleles^13^.

To bypass the requirement for known HLA-peptide combinations, peptide pools or whole pathogens can be used for short (6-24h) *in vitro* stimulation, followed by isolation and single-cell sequencing of T cells by proxy markers of antigen-specificity. Activated T cells can be identified by the expression of activation-induced markers (AIM) on their cell surface^14,15,16^, and AIM+ cells can be sorted for sequencing to determine their TCR sequences^17,18,19,20^. This approach was successfully used to discover novel features of Hepatitis C virus-specific CD4 T cells^20^, SARS-Cov2-specific CD4 and CD8 T cells^17,18^, and cow milk allergy-specific CD4 T cells^19^. One downside of this method is that the combination of activation markers for identifying antigen-specific T cells varies between pathogens, tissues, and length of stimulation and thus requires careful optimization^16,20^. Additionally, the identified cells by definition are in an activated state, which may not reflect their natural state *in vivo*. Finally, both AIM assays and HLA-peptide sorting based approaches require large volumes of blood to obtain sufficient numbers of antigen specific T cells, which typically have frequencies of less than 1 in 10,000.

We and others have used *in vitro* stimulation for ∼14 days of peripheral blood mononuclear cells (PBMC) with peptides to expand antigen-specific T cells in the context of infection, vaccination, and allergy^21,22,23,24,25,26,27^. This is particularly useful for samples with rare/low frequency of antigen-specific T cells, e.g., self-antigen-specific T cells in the case of Parkinson’s disease^27,28,29^. Another advantage of this method is that, following expansion, the antigen-specific T cells can be directly studied, without the need for further isolation^26,27^. However, following a 14-day expansion, the gene expression phenotype of these cells will be quite different from their *in vivo* state.

Here, we combined our previous experience with *in vitro* expansion of antigen-specific T cells using peptide pools to develop a protocol to identify antigen-specific TCRs called **<u>P</u>**eptide-**<u>D</u>**riven **<u>I</u>**dentification of **<u>TCR</u>**s (PDI-TCR). In PDI-TCR, PBMC are stimulated with a peptide pool of interest to expand antigen-specific T cells, and the TCR repertoire of the expanded cells is determined 14-days post-stimulation by bulk TCR β sequencing. The same process is repeated with a non-overlapping control peptide pool, and a statistical analysis is performed to identify antigen-specific TCRs (i.e., significantly expanded with the peptide pool of interest) versus unspecific expanded TCRs (i.e., expanded in both conditions). The identified TCRs are then used to examine the direct *ex vivo* phenotype of antigen specific T cells.

We applied PDI-TCR to study *Mycobacterium tuberculosis (Mtb)*-specific cells in a cohort of patients with active tuberculosis (TB) and Mtb-sensitized healthy individuals (asymptomatic, but with Mtb immune memory, as demonstrated by a positive interferon-gamma release assay, IGRA+). In combination with single-cell mRNA and TCR sequencing of *ex vivo* sorted CD4 T cells, we successfully determined the frequency and transcriptome of Mtb-specific T cells in TB and IGRA+ cohorts, as well as through anti-TB therapy, highlighting novel phenotypes and dynamics of antigen-specific T cells throughout TB disease and treatment. We also identified a population of ‘unspecific expanded’ T cells that showed a distinct TCR clonality and gene expression profile.

## Methods

### Ethics statement

Human study participants were enrolled at the National Hospital for Respiratory Diseases, Welisara (Sri Lanka), or The South African Tuberculosis Vaccine Initiative, Western Cape Province (South Africa). Ethical approval to carry out this work was maintained through the La Jolla Institute for Immunology Institutional Review Board (IRB) or the Human Research Ethics Committee of the University of Cape Town. The University of Colombo Ethics Review Committee is the National Institute of Health registered IRB for Kotelawala Defence University, Sri Lanka. All participants provided written informed consent before participation in the study. All samples were obtained for specific use in this study.

### Participants and samples

Tuberculosis (TB) was defined as 1) the presence of clinical symptoms and/or radiological/histological evidence of pulmonary TB and 2) microbiologically confirmed (by sputum or Mtb-culture), in the absence of any other significant comorbidity-morbidity. Blood samples were obtained at diagnosis, mid-treatment, end of treatment and one year post diagnosis. Anti-TB therapy was a standard regimen for drug-susceptible *Mtb* consisting of an intensive phase of two months with isoniazid (INH), rifampin (RIF), pyrazinamide (PZA), and ethambutol (EMB) followed by a continuation phase of four months with INH and RIF^30^. IGRA+ controls were healthy individuals with no symptoms or history of TB, with Mtb-specific immune memory confirmed by a positive IFNγ release assay (IGRA) at the time of recruitment. IGRA-healthy controls were healthy individuals with no Mtb-specific immune memory as confirmed by a negative IGRA assay. All participants were confirmed HIV-negative. Participants were bled by venipuncture for a small blood draw (10-20 ml). PBMC were obtained by density gradient centrifugation (Ficoll-Hypaque, Amersham Biosciences/ GE Healthcare) from whole-blood samples, according to the manufacturer’s instructions. Cells were resuspended in FBS (Gemini Bio-Products) containing 10% DMSO (Sigma) and cryopreserved in liquid nitrogen.

### PBMC thawing

Cryopreserved PBMC were quickly thawed by placing each cryovial in a 37 °C water bath for 2 min, and cells transferred into 9 ml of cold medium (RPMI 1640 with L-Glutamin and 25 mM Hepes (Omega Scientific), supplemented with 5% human AB serum (GemCell), 1% Penicillin Streptomycin (Gibco) and 1% Glutamax (Gibco) and 20 U/mL Benzonase Nuclease (Millipore). Cells were centrifuged and resuspended in media to determine cell concentration and viability using Trypan blue and a hematocytometer. Cells were then kept at 4 °C until used for flow cytometry or cell sorting.

### Peptide pools

For the expansion of Mtb-specific T cells we used the MTB300 peptide pool, which is composed of 300 Mtb-derived immunodominant CD4 T cell epitopes^31^. For the expansion of *Bordetella pertussis* (BP)-specific T cells we used the BP(E)VAC peptide pool, which is composed of 132 peptides with experimentally validated immunodominant CD4 T cell epitopes against the 4 acellular *Bordetella pertussis* vaccine antigens FHA, FIM2/3, PRN, and PtTox^32^. To create the negative megapool (NMP), an initial set of 100,000 unique, random peptides were generated in R. PepSySco was then used to predict the likelihood that the peptides in the pool can be synthesized successfully. The top 1,000 peptides from PepSySco (score of 0.92 and up) were then used to run BLAST against the NCBI nonredundant protein (nr) database. 529 peptides had hits from BLAST and were thus removed from the peptide set. The remaining 471 peptides were then run in PEPMatch to confirm the peptides do not have any sequence similarity to proteins from the archaea, bacteria, eukaryote, or virus proteomes. Finally, the 7-allele method in TepiTool was used to generate MHC class II prediction for the 471 peptides. From the TepiTool output, the top 100 scoring peptides were chosen to generate the NMP.

### Antigen-specific T cell expansion

PBMC were plated at 2.5×10^5^ cells/well (for TB patients) or 5.0 ×10^5^ cells/well (for IGRA+ individuals) in a final volume of 250 µl of media (RPMI 1640 with L-Glutamin and 25 mM Hepes (Omega Scientific), supplemented with 5% human AB serum (GemCell), 1% Penicillin Streptomycin (Gibco) and 1% Glutamax (Gibco)) per well (corresponding to 1×10^6^ cells/mL final concentration) in a 96-well U bottom plate. Cells were stimulated with either Mtb-specific (i.e., MTB300) or BP-specific (i.e., BP(E)VAC) peptide pools at 2 μg/mL final concentration and incubated at 37 °C for 14 days. From each well, 125 μl of the culture supernatant (corresponding to half of the total culture volume) was replaced with fresh media every three to four days and was supplemented with 0.02 U/μL of IL-2 (Prospec) on days 4, 8, and 12. After 14 days, cells were spun before proceeding with DNA extraction.

### Ex vivo CD4 T cell isolation

After PBMC thawing, CD4 T cells were purified using the EasySep™ Human CD4 Positive Selection Kit II (STEM cell^TM^ Technologies), according to the manufacturer’s instructions. Approximately 2 million PBMC were used for CD4 T cell isolation.

### Bulk β TCR sequencing

DNA extraction (DNeasy Blood and Tissue kit; Qiagen) was performed according to the manufacturer’s instructions for both *ex vivo* purified CD4 T cells and 14-day cultured PBMC. TCR sequencing of the V-β chain was performed by Adaptive Biotechnologies and their ImmunoSeq platform. *Ex vivo* CD4 T cells were sequenced on “deep” mode while 14-day cultured PBMC were sequenced on “survey” mode. *Ex vivo* CD4 T cells were sequenced at a higher depth than 14-day cultured PBMC to increase the likelihood of detecting rare clonotypes and get a more accurate account of the global TCR repertoire diversity in each sample.

### Bulk β TCR sequencing analysis

All statistical analyses were performed on R version 4.4.0. To determine the abundance of a TCR β clonotype (defined as one unique productive CDR3 amino acid sequence), the number of occurrences, or “templates”, for each TCR β clonotype was quantified per sample. Since TCR sequencing was performed on genomic DNA, and most T cells are expected to contain one single rearranged CDR3 DNA sequence per cell^33,34^, we assumed that one occurrence (or template) equals one cell. TCR β clonotypes were ranked based on total abundance, defined as the sum of abundance of a TCR β clonotype in Mtb- and BP-stimulated samples per donor (or per replicate, if more than one culture per peptide pool were performed). The top 1% most abundant clonotypes per donor were selected to perform a Fisher’s exact test comparing the abundance between Mtb- and BP-stimulated samples. A pseudo count of 1 was added to each abundance to correct for null values, before calculating fold changes. Fold changes were calculated as the ratio of the abundance of a clonotype in Mtb-versus BP-stimulated samples.

### Single-cell RNA and TCR sequencing using 10X

#### Flow cytometry and TotalSeq^TM^-C antibody staining

Freshly thawed PBMC were incubated for 10 minutes with 1 µl of UV Zombie Fixable Viability Kit (BioLegend) at 1:100 dilution in 1X PBS following the manufacturer’s protocol. To neutralize unbound viability dye, 10% Fetal Bovine Serum (FBS) in 1X PBS was added to the cells followed by a centrifugation step. FcR blocking reagent (BioLegend) at 1:50 dilution from the stock reagent solution was added to the cells and samples were further incubated for 10 minutes at room temperature. For oligonucleotide-conjugated antibody staining, 2 μl of each TotalSeq^TM^-C antibodies (**Table S1**) were added per sample in a final staining volume not exceeding 100 µl in 1X PBS and cells were incubated for 15 minutes at room temperature. All samples received tagging with ADT-CD4 and ADT-HLA-DR TotalSeq^TM^-C antibodies, while the remaining TotalSeq^TM^-C antibodies were used as sample barcodes (i.e., only one antibody per sample). After staining, cells were spun down, and excess unbound antibody was removed. A cocktail of fluorochrome-conjugated antibodies (**Table S2**) was added to each sample, along with 10 µl BD Horizon TM Brilliant stain buffer plus (BD Biosciences), and PBS for a final volume of 150 µl. Cells were incubated for 20 minutes in the dark at room temperature. Cells were washed twice in 1X PBS to remove all unbound antibodies and stored at 4 °C protected from light for up to 4 hours until flow cytometry acquisition. Live singlet CD3+ CD19-CD14-CD4+ cells that are HLA-DR+/ HLA-DR-were sorted using a BD Symphony S6 sorter (see gating strategy **Figure S2A**). A target of 3,333 cells per sample per population were set for sorting cells for sequencing. However, the actual cell numbers ranged between 241-3,333 depending on the cell viability of each sample.

#### 5’ single-cell RNA-sequencing

Cells were resuspended in ice-cold PBS with 0.04% BSA in a final concentration of 1,800 cells/μL. Single-cell suspensions were then immediately loaded on the 10X Genomics Chromium Controller with a loading target of 30,000-60,000 cells. Libraries were generated using the Chromium Next GEM Single Cell 5′ Reagent Kit v2 (Dual Index) per the manufacturer’s instructions, with additional steps for the amplification of HTO barcodes and V(D)J libraries (10X Genomics). Libraries were sequenced on Illumina NovaSeq with a sequencing target of 30,000 reads per cell RNA library, 5,000 reads per cell HTO barcode library, and 5,000 reads per cell for V(D)J libraries.

#### Bioinformatic analysis

Raw reads were aligned and quantified using the Cell Ranger Multi pipeline (version 5.0.0) against the GRCh38 (2020-A) human reference genome and the VDJ reads were aligned with refdata-cellranger-vdj-GRCh38-alts-ensembl-5.0.0. GEX and HTO data were aggregated as feature/barcode matrices and imported into the R environment using Seurat where additional QCs and downstream analysis were conducted. Before creating the Seurat object, in the GEX matrix, TCR genes belonging to each category of TCR α, β, γ and δ were averaged for each cell to prevent the formation of individual TCR gene expression clusters in downstream analysis. The resulting gene expression matrix was normalized with ‘SCTransform’, followed by Principal Component Analysis (PCA) and clustering with ‘FindNeighbors’ and ‘FindClusters’. Dimensionality reduction was achieved by UMAP. The ‘FindAllMarkers’ function was used to identify the characteristic genes for each cluster with the MAST test. These two processes were iterated upon, varying the QC thresholds and clustering parameters until the identified clusters were stable, according to a bootstrap analysis. Further selection for CD4+ T cells was done using ADT-CD4 expression. Cells with no ADT-CD4 expression were excluded and the new object was re-transformed. For TCR analysis, filtered contigs from Cell Ranger ‘multi’ results were imported into R and processed to assign clonotypes. A clonotype was defined by the TRA and TRB combination. Clonotype sizes were categorized based on their size at each cohort and visit: singletons (clone size of 1), small (1 < X <= 5), medium (5 < X <= 20) and large (20 < X <= 100) after down sampling each cluster to represent the same number of cells per cohort and visit. Mtb-specific and unspecific TCRs identified with PDI-TCR were mapped to the single-cell 10X dataset by matching amino acids in the cdr3 region. Only cells with a paired α and β sequence were considered for downstream analysis of antigen-specific TCRs.

TCR clonotypes with paired α and β chains were selected, and the use of TRA and TRB genes by each distinct TCR was quantified for each donor. The count of each TRA or TRB gene use across the donors was summed. Gene frequencies were determined by dividing the sum of each TRA or TRB gene by the sum of all TRA or TRB genes. A fold change between the Mtb- and unspecific frequencies was calculated, and only those with a fold change >2 between Mtb versus other or unspecific versus other were considered preferentially used.

#### Statistical Analysis

All statistical analyses were performed on R version 4.4.0 using R packages: Tidyverse, dplyr and ggplot2 or GraphPad Prism version 10.2.3 (347). We performed Mann-Whitney U test on unpaired samples using the ggpubr package, and Wilcoxon signed-rank test on paired samples using the robustrank package. Frequency of cells in the HLA-DR+ clusters (cluster 1, 4, 6 and 7) were adjusted to correct for the frequencies of HLA-DR+ cells in total CD4 T cells in each sample, as determined by flow cytometry. Two donors had missing paired visits and were thus excluded from down-stream analysis.

### TCR validation experiments

#### Generation of TCR retroviruses

For TCR validation, we selected a paired Mtb-specific sequence with the highest number of Mtb-specific cells at diagnosis (3 cells) in the 10X dataset and had PBMCs left for subsequent testing if required. We used the amino acid sequence of the β and α CDR3 chains, and their gene usage to generate the full-length nucleotide sequences of the TCR α and β chains by Stitchr^35^. The full-length TCR sequence was generated by switching the human constant regions with murine TCR β and α constant regions and placing a Furin-P2A sequence between the β and α sequence as described previously^36^. The TCR sequence was codon optimized for expression in human cells using NovoPro ExpOptimizer and was synthesized as a gBLOCK by IDT. The TCR gBLOCK was inserted into a MSGV1 retrovirus backbone (Addgene plasmid #122728) via gibson assembly. TCR retroviral supernatant was generated by transfection of 293GP cells with the TCR-containing MSGV1 DNA along with envelope protein RD114a (Addgene plasmid #17576). 48 hours post transfection, TCR retroviral supernatant was harvested and snap-frozen.

#### Cell Transduction

To generate TCR-cell lines, retroviral supernatant was used to transduce triple-reporter jurkats engineered to express NFAT-eGFP, NF-κB-CFP and AP-1-mCherry upon T cell activation^37^. Jurkats were plated at 3.75 × 10^6^ cells per well in a 24 well plate (GenClone #25-107) in 1X RPMI medium (Fisher Scientific #11-875-093) supplemented with 10% Fetal Bovine Serum (Gemini Bio Products #100-106-500) and 1% Penicillin: Streptomycin (Gemini Bio Products #400-109). 300 IU/mL IL-2 (Prospec Bio #Cyt209) and 50 ng/mL anti-human CD3 (clone OKT3, Invitrogen) were added to the culture and the cells were rested for 48 hours. Non-tissue culture 24-well plates (USA Scientific #5665-5185) were coated with RetroNectin (Takara Bio T100A) at 500 ul/well and incubated at 4 °C until the jurkats were ready for transduction. After 48 hours, the cells were transduced with retroviral supernatant for 72 hours via spinfection as described previously^38^. Briefly, retronectin-coated plates were spun at 2000 RPM at 32 °C for 1 hour and then blocked with 2% BSA (Millipore Sigma #A9418) in PBS for 30 minutes. Retroviral supernatant was added to the plates and spun at 2000 RPM for 2 hours at 32 °C. Jurkats were then added to the plates at 250,000 cells per well in cell media supplemented with IL-2. The plates were spun a final time at 1500 RPM for 10 minutes and then placed in an incubator overnight. The next day, transduced cells were transferred to tissue-culture coated 24-well plates and left to incubate until 72 hours post-transduction.

#### TCR-cell line and lymphoblastoid cell line (LCL) coculture

A lymphoblastoid cell line (LCL) was generated by Epstein-Barr virus transformation^39^. Briefly, PBMC isolated from the subject were cultured with EBV (ATCC #VR-1492) for several weeks to generate a continuously growing antigen-presenting cell line. To assess TCR reactivity, TCR-jurkats were cocultured with LCLs pulsed with MTB300 and NMP pools. LCLs were plated at 500,000 cells/ml in 1X RPMI supplemented with 10% Fetal Bovine Serum, 1 % non-essential amino acids (Thermo Fisher #11140050), and 1% Penicillin: Streptomycin. LCLs were pulsed with 0.1 uM, 1 uM, and 10 uM of peptide pools and incubated at 32 °C overnight. Un-pulsed co-cultures contained only the TCR-jurkats and LCLs. The next day, LCLs were spun down at 500 g for 5 minutes and washed twice with PBS. The cells were resuspended in cell media and 50,000 TCR-jurkats were added to the wells and incubated for 6 hours. T cell-jurkat activation was assessed via flow cytometry.

#### Flow cytometry

For the TCR-Jurkat experiments, the cells were spun down and incubated with 10% FBS in PBS for 10 min at room temperature. Cells were then stained with viability dye and surface antibodies for 20 minutes at room temperature. The cells were then spun and washed twice with PBS and then resuspended in PBS supplemented with 2 mM EDTA and 0.5% BSA (Sigma #A-3294) and acquired on a BD LSRII. Data was analyzed on FlowJo (version 10.10) and activated cells were defined as live, singlet, TCR+ cells expressing either NFAT-eGFP, NF-κB-CFP and AP-1-mCherry.

## Results

### Development of the PDI-TCR method on a pilot cohort of 4 TB patients

The PDI-TCR experimental and analytical framework is presented in **Figure 1A** & **Figure 1B**. We first tested the method on a pilot cohort of four TB patients with PBMC collected at diagnosis (see **Table 1** for patient demographics). PBMC were stimulated with Mtb-specific or an unrelated BP-specific peptide pools for 14 days to expand Mtb-specific or BP-specific CD4 T cells, respectively (see methods for details). We have previously shown that Mtb-specific T cells can be detected using the Mtb-peptide pool in TB individuals^40,41^, and BP pool reactivity has been found in healthy adults based on childhood *B. pertussis* vaccination^32,42,43,44,45^. To estimate the technical variability of the assay, we included two replicates for each stimulation condition for each patient.

**Figure 1:**
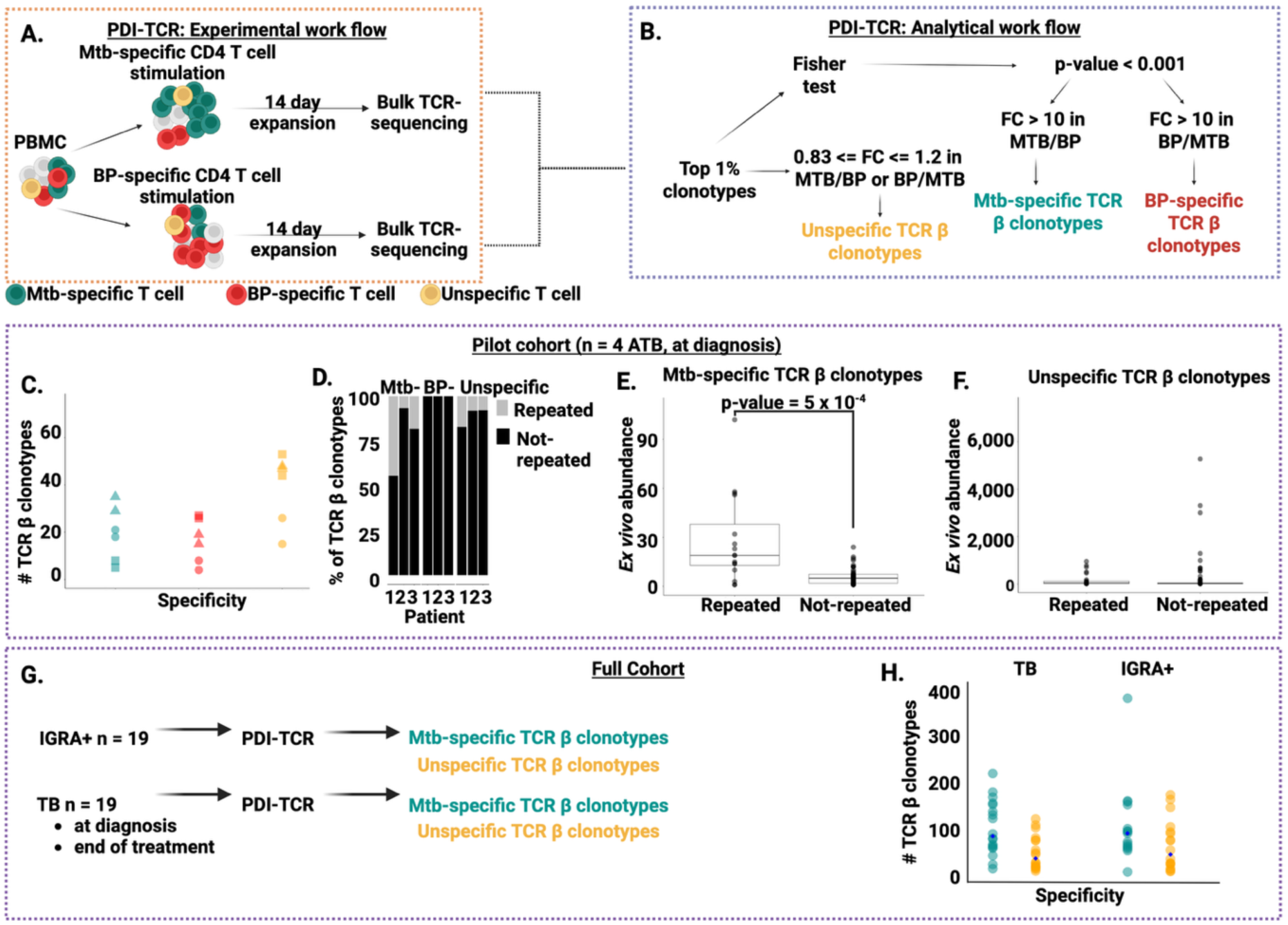
Comprehensive Analysis of PDI-TCR Workflow, Clonotype Specificity, and Abundance in Mtb-vs BP-Cultures. A) Experimental workflow of PDI-TCR. Participants with TB (n=4) were recruited for blood donation at the time of diagnosis. PBMC were cultured in the presence of MTB300 or BP(E)VAC peptide pools. Cells were harvested after 14 days for bulk survey TCR β sequencing. B) Analytical workflow of PDI-TCR. TCR β clonotypes were ranked by their total abundance in Mtb- and BP-culture, and their top 1% were selected for the Fisher’s exact test. TCR β clonotypes were categorized as Mtb-, BP-, or unspecific based on the defined statistical criteria. Mtb- and BP-specific TCR β clonotypes had a p-value < 0.001 and an FC > 10. unspecific TCR β clonotypes had a FC between 0.83 and 1.2. C) Total number of Mtb-, BP-specific and unspecific TCR β clonotypes that were identified with PDI-TCR in each sample (n = 6). Dark cyan: Mtb-specific TCR β clonotypes; Red : BP-specific TCR β clonotypes; Yellow: Unspecific TCR β clonotypes. D) Proportion of TCR β clonotypes found to be repeated and not repeated between technical replicates in each patient (n = 3). Grey: Repeated TCR β clonotypes; Black: Not repeated TCR β clonotypes. Only a small proportion of TCR β clonotypes were found to be repeated. E) *Ex vivo* abundance of the repeated versus not repeated Mtb-specific TCR β clonotypes in each patient (n = 3). Each dot represents a TCR β clonotype plotted by its *ex vivo* abundance (y-axis). Repeated Mtb-specific TCR β clonotypes had a significantly higher *ex vivo* abundance compared to their non-repeated counterparts. F) *Ex vivo* abundance of the repeated versus not repeated unspecific TCR β clonotypes in each patient (n = 3). Each dot represents a TCR β clonotype plotted by their *ex vivo* abundance (y-axis). Repeated unspecific TCR β clonotypes did not have a higher *ex vivo* abundance compared to their not repeated counterparts. G) This figure illustrates the workflow of the PDI-TCR analysis conducted on the full cohort. The cohort consisted of a total of 19 IGRA+ individuals and 19 TB individuals. Each TB individual had a sample from a matched visit at the time of diagnosis and at the end of treatment. The PDI-TCR was performed on the samples to identify Mtb- and unspecific TCR β clonotypes for each donor. H) Total number of Mtb- and unspecific TCR β clonotypes that were identified with PDI-TCR for each donor (n = 38). Dark cyan: Mtb-specific TCR β clonotypes; Yellow: Unspecific TCR β clonotypes.

**Table 1:** Patient demographic data.

|  | Pilot (n=4) | Full Cohort (n=58) |  |  |
| --- | --- | --- | --- | --- |
|  | TB (n = 4) | TB (n= 19) | IGRA+<br>(n=20) | IGRA- (n=20) |
| Age, years<br>(median [IQR]) | 49 (22-55) | 36 (25-48) | 38 (29-42) | 37 (30-44) |
| Female (%) | 25% | 47% | 50% | 50% |
| Male (%) | 75% | 53% | 50% | 50% |

After 14 days, the TCR β repertoire of all cells remaining in culture was determined by bulk TCR-sequencing. **Table S3** summarizes the approximate number of cells expressing a given TCR β, unique TCR β clonotypes, and TCR β clonotype singletons (i.e., with an abundance of 1 cell) retrieved in each sample. The number of unique TCR β clonotypes in each patient ranged between 27,231 to 59,912 for *ex vivo* samples, and between 5,659 to 16,176 for *in vitro* stimulated samples (**Table S3**). For both *ex vivo* and *in vitro* stimulated samples, singletons represented the majority of TCR β clonotypes retrieved (80-92%, **Table S3**).

To identify TCR β clonotypes that were significantly expanded after Mtb- or BP-peptide pool stimulation, we applied two criteria. First, a Fisher’s exact test that compares pre-vs. post culture frequency of a given clonotype with an un-adjusted p-value cutoff <0.001 to identify clonotypes with a *statistically significant increase* after culture. Second, a simple ‘top 1%’ cutoff for clonotypes after the 14 day culture that measures *high abundance* after culture. While the statistical increase criterion is more precise, the high abundance criterion has the advantage of only needing the post-expansion sample. For the Mtb-cultures, we found that TCR β clonotypes that were statistically increased after culture were also in high abundance (**Figure S1A** and **S1B**). In contrast, TCR β clonotypes that were statistically increased in BP-cultures reached the high abundance threshold less frequently. This likely reflects that some of the TCRs marked as statistical expanded for BP are false-positives given the large number of TCRs tested and that we are expecting much fewer antigen-specific T cells for BP compared to Mtb in TB patients, where the exposure is due to vaccinations that might be decades ago, as opposed to active Mtb infection.

Surprisingly, we also found a population of TCR β clonotypes that were statistically increased after culture with either Mtb or BP peptides, even though the peptides had no sequence overlap (**Figure S1B**). These TCR β clonotypes were also found in high abundance after both the Mtb- and BP-cultures (**Figure S1C**) and could have easily been misidentified as being Mtb- or BP-specific. This highlights the importance of doing additional stimulations other than the one of interest to identify truly antigen-specific clonotypes.

Next, we wanted to develop a scheme that does not require *ex vivo* sequencing, but only the Mtb- and BP-expanded cultures. We restricted our analysis to the top 1% high abundance TCR β clonotypes after each culture, which reduces the number of clonotypes under consideration, and reduces multiple hypothesis testing. As shown in **Table S4**, this reduces the number of clonotypes considered to <300 per sample. Next, we applied the statistical increase test, but now comparing expansion in Mtb-vs. BP-cultures. A p-value of <0.001 reduced the FDR to less than 1% and retained >30 significant TCR β clonotypes (33-78, **Table S4**) in 3 out of 4 patients.

Patient 4, whose samples showed the highest number of singletons and the least number of significantly expanded Mtb-TCR β clonotypes (**Table S3** and **Figure S1B**), had very few significant TCR β clonotypes across both replicates regardless of the p-value used (**Table S4**). This patient was thus excluded from downstream analysis, as its expansion cultures seemed to have failed.

Inspecting the clonotypes marked as Mtb-expanded or BP-expanded using the abundance and statistical criteria showed that some had very high cell numbers in both Mtb and BP-cultures, with differences in abundance lower than 20%. While the difference in these cell numbers might be statistically significant, our opinion is that it would not be biologically correct to call these clonotypes solely Mtb-specific or BP-specific. Thus, we introduced a *fold change* cutoff > 10 as an additional filter to select significant TCR β clonotypes with at least 10-fold disparity in abundance between Mtb- and BP-stimulated cultures.

Finally, we wanted to follow up the surprising finding of clonotypes expanded similarly with different peptide pools. We identify these unspecific TCR β clonotypes by using the top 1% abundance threshold and in addition asking for a limited fold-change between the Mtb and BP cultures between 0.83 and 1.2. The final PDI-TCR analysis workflow is presented in **Figure 1A** & **Figure 1B**. To summarize, <u>antigen specific</u> TCR β clonotypes are defined by (1) an *abundance* cutoff of top 1%, combined with a (2) *statistical* cutoff for a Fishers test p < 0.001, and (3) a *fold change* cutoff > 10. In contrast, <u>unspecific</u> TCR β clonotypes are those within the top 1% abundance after culture that have a FC cutoff between 0.83 and 1.2.

The number of Mtb-specific, BP-specific, or unspecific TCR β clonotypes obtained per sample in the pilot cohort is summarized in **Figure 1C** and further illustrated in **Figure S1D** for each patient and replicate. The highest number of TCR β clonotypes was found for the unspecific category (15-53), followed by Mtb-specific (5-35), and BP-specific (4-27) (**Figure 1C**). For each patient, numbers were highly similar between replicates (**Figure 1C**). Thus, combining an experimental and a statistical workflow, PDI-TCR can be used to identify antigen-specific and unspecific TCR β clonotypes from PBMC samples.

### Ex vivo abundance determines PDI-TCR sensitivity to antigen-specific TCR β clonotypes

Next, we tested the sensitivity of the PDI-TCR assay by comparing the Mtb-specific, BP-specific and unspecific TCR β clonotypes found in the different replicates of each patient. A TCR β clonotype found in both replicates was labeled as “repeated”, while a TCR β clonotype present in only one replicate was labelled as “not repeated”. Only a small fraction of TCR β clonotypes were repeated, ranging from 7-44% for Mtb-specific TCR β clonotypes, 8-17% for unspecific TCR β clonotypes, and 0% for BP-specific TCR β clonotypes (**Figure 1D**). We hypothesized that the likelihood of observing an expanded clone might be driven by its frequency in the initial sample. Thus, we compared the *ex vivo* abundance of repeated and non-repeated TCR β clonotypes. Indeed, repeated Mtb-specific TCR β clonotypes had a higher *ex vivo* abundance compared to their non-repeated counterparts (**Figure 1E**).

For unspecific TCR β clonotypes, which we defined as having high and similar abundance in both Mtb- and BP-cultures, we found no correlation between their *ex vivo* frequency and their repeatability (**Figure 1F**). This is no surprise as the majority of unspecific TCR β clonotypes (39%-94%) showed decreasing abundance between *ex vivo* and post-stimulation conditions (**Figure S1E**). This suggest that the majority of the unspecific TCR β clonotypes do not expand in response to the stimulation but rather remain in culture at similar or slightly decreased frequency.

### Identification of Mtb-specific and unspecific TCR β clonotypes in a larger cohort of TB and Mtb-sensitized individuals using PDI-TCR

Next, we tested the utility of the PDI-TCR method by applying it to a larger cohort. We used PBMC samples from 19 TB patients collected at diagnosis, as well as 19 healthy individuals with no symptoms or history of TB with Mtb-specific immune memory defined as a positive IFNγ release assay (Mtb-sensitized, or IGRA+ cohort) (see **Table 1** for cohort demographics). Most TB patients have lymphopenia at the acute phase of infection, resulting in a limited number of PBMC retrieved in the diagnosis blood sample. Thus, to increase the number of detected TCR β clonotypes for each TB patient, we applied PDI-TCR on an additional longitudinal PBMC sample collected after completion of six-month anti-TB therapy (end-treatment sample). We applied the experimental and analysis workflows of PDI-TCR presented in **Figure 1A** and **1B** to identify Mtb-specific and unspecific TCR β clonotypes for each sample in the full cohort (**Figure 1G**). While there was variability between participants, the median number of Mtb-specific and unspecific TCR β clonotypes was similar between TB and IGRA+ individuals and between categories (78 in TB and 84 in IGRA+ Mtb-specific TCR β clonotypes and 30 in TB and 38.5 in IGRA+ unspecific TCR β clonotypes, **Figure 1H**). Thus, using the PDI-TCR assay, we successfully identified hundreds of Mtb-specific and unspecific TCR β clonotypes in TB patients and Mtb-sensitized (IGRA+) individuals.

### Ex vivo single-cell transcriptomic and TCR analysis of Mtb-specific and unspecific CD4 T cells

To characterize the phenotype of cells with Mtb-specific and unspecific TCRs as identified with PDI-TCR in the full cohort, we interrogated the single-cell transcriptome and TCR repertoire of *ex vivo* CD4 T cells in the same PBMC samples (**Figure 2A**). For the TB cohort, we included two additional longitudinal PBMC samples: three months after initiation of anti-TB therapy (mid-treatment), and six months after completion of anti-TB therapy (1 year post-diagnosis), for a total of four samples per patient with TB. Additionally, we included a cohort of 20 IGRA-healthy individuals as controls (**Table 1**). We previously found that at diagnosis of TB, HLA-DR is a marker for Mtb-specific CD4 T cells^40^. Since HLA-DR positive cells typically represent less than 10% of total CD4 T cells, we sorted an equal number of HLA-DR+ CD4 T cells and HLA-DR-CD4 T cells (see gating strategy in **Figure S2A**) and pooled them together at 1:1 ratio for single-cell RNA and TCR sequencing to enrich the sequenced sample for likely antigen specific cells.

**Figure 2:**
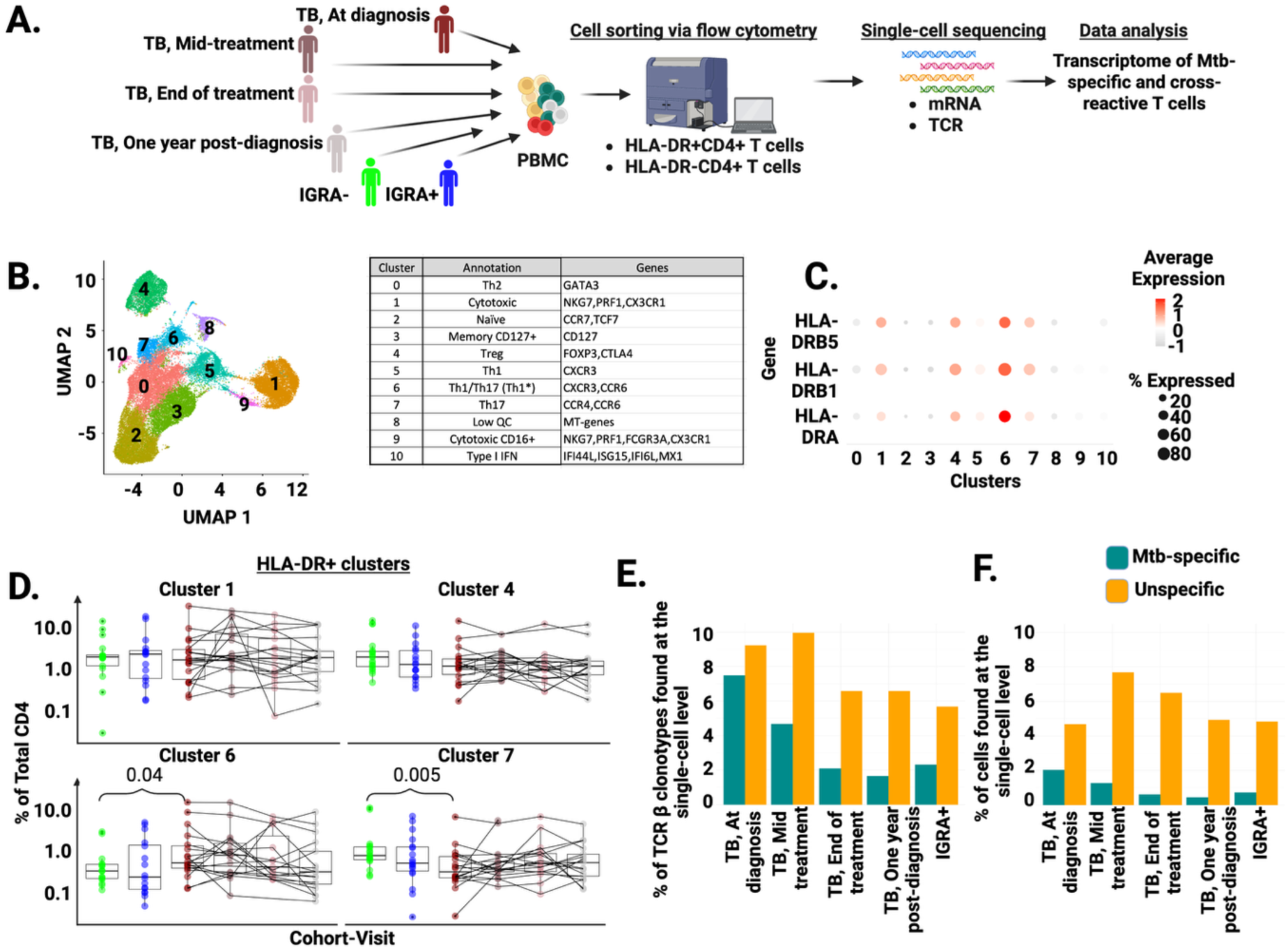
Mapping PDI-TCR identified TCRs at the single-cell level. A) Experimental design. PBMCs from the same ATB patients (n = 19) and their matched visits (at diagnosis, mid-treatment, end of treatment and one-year post-diagnosis), IGRA+ (n = 20) and IGRA-(n = 20) individuals were sorted *ex vivo* for HLA-DR+ and HLA-DR-CD4 T cells in equal numbers and were subjected to single-cell 5’ mRNA and TCR-sequencing. Downstream single-cell analysis was performed. The PDI-TCR identified Mtb- and unspecific TCR β clonotypes from each TB and IGRA+ individual was mapped at the single-cell level by matching the β CDR3 sequences. B) Based on RNA profiles of cells from all samples that were sequenced (n = 116), 11 RNA clusters were identified and annotated based on the expression of canonical markers of cell types. C) Dot plot of the three HLA-DR genes-HLA-DRB5, HLA-DRB1, HLA-DRA. The size of each dot is the fraction of cells expressing the feature, the color scale represents the expression level. Clusters 1, 4, 6 and 7 were all HLA-DR+. D) Adjusted HLA-DR+ Cell Counts in CD4 T Cell Clusters. This figure shows the adjusted HLA-DR+ cell counts for clusters 1, 4, 6, and 7. The number of cells in each cluster was adjusted to reflect the true frequency of HLA-DR+ cells in total CD4 T cells (y-axis) within each sample per donor (n=57). Cluster 6 exhibited the highest frequency of HLA-DR+ CD4 T cells in TB patients at diagnosis, compared to the respective treatment points and healthy individuals. Dark red: TB at diagnosis; Dark pink: TB mid-treatment; Light pink: TB end of treatment; Grey: One-year post diagnosis; Blue: IGRA+; Green: IGRA-. E) *Ex Vivo* Mapping of PDI-TCR identified Mtb- and unspecific TCR β clonotypes. This figure shows the percentage of PDI-TCR identified Mtb- and unspecific TCR β clonotypes found directly *ex vivo* at the single cell level (y-axis) in TB (n=16) and IGRA+ (n=19) individuals. The highest proportion of Mtb-specific TCR β clonotypes in TB patients were found at diagnosis sample *ex vivo*, whereas majority of unspecific TCR β clonotypes were found *ex vivo* at the TB mid-treatment samples. Dark cyan bars: Mtb-specific TCR β clonotypes; Yellow bars: Unspecific TCR β clonotypes. F) Proportion of cells of Mtb- and unspecific TCR β clonotypes found *ex vivo* at the single-cell level. This figure shows the percentage of Mtb- and unspecific cells found directly *ex vivo* as a fraction of total cells found at each cohort and visit (y-axis). Highest proportion of Mtb-specific cells were found at TB at diagnosis. Highest proportion of unspecific cells were found at TB mid-treatment.

UMAP and clustering analysis on the mRNA data identified eleven clusters (**Figure 2B**). Each cluster was manually annotated (**Figure 2B**) using their top expressed genes (**Figure S2B** and **Table S5**) and known chemokine receptors for T helper subsets (**Figure S2C**). We identified naïve T cells (cluster 2), Tregs (cluster 4), cytotoxic T cells (clusters 1 and 9), Th1 (cluster 5), Th2 (cluster 0), Th17 (cluster 7) and Th1* (also referred to as Th1/Th17, cluster 6). There was also one cluster of undefined memory CD127+ T cells (cluster 3), and one cluster with high expression of type 1 IFN signaling genes (cluster 10). Cluster 8 presented high expression of mitochondrial genes and was thus annotated as low quality. Four clusters showed positive expression of HLA-DR: clusters 1, 4, 6 and 7 (**Figure 2C**). Cluster 6 (Th1*), was the cluster with the highest HLA-DR expression.

We compared the CD4 T cell cluster composition across different cohorts by taking in consideration whether the cluster was classified as HLA-DR positive or negative (**Figure 2C**) and the frequency of HLA-DR+ cells within total CD4 T cells determined by flow cytometry (**Figure S2D**). This was necessary due to the enrichment for HLA-DR+ cells in the sorted CD4 T cell population that was used for single-cell sequencing. Out of the 19 TB patients subjected to single-cell TCR sequencing, two participants had missing paired visits and were thus excluded.

Among the four HLA-DR+ clusters, only cluster 6 (Th1*) showed significant higher frequency in TB at diagnosis compared to the IGRA-cohort (**Figure 2D**). Conversely, cluster 7 (Th17) showed a significantly decreased frequency in TB at diagnosis compared to the IGRA-cohort (**Figure 2D**). Within the HLA-DR-clusters, there were no differences between TB diagnosis, IGRA+ and IGRA-cohorts (**Figure S2E**). Cluster 3 (memory CD127+) showed an increased frequency in TB from diagnosis to mid-treatment, while cluster 9 (cytotoxic CD16+ T cells) showed a decreased frequency in TB at mid-treatment compared to the one-year follow up visit (**Figure S2E**).

Next, we determined the number of Mtb- and unspecific TCR clonotypes identified with PDI-TCR that were found *ex vivo* using the single-cell TCR sequencing data. We found that 16 TB patients and 19 IGRA+ individuals had matched cells based on the CDR3 amino acid sequences of the TCR β clonotypes. A total of 1390 & 1512 Mtb-specific TCR clonotypes were identified by PDI-TCR combining TB and IGRA+ individuals, respectively. In addition, a total of 683 & 1039 unspecific TCR clonotypes were identified in all combined TB and IGRA+ individuals. We calculated the proportion of Mtb-specific TCR clonotypes found *ex vivo* as a fraction of the total number of Mtb-specific TCR β clonotypes identified by PDI-TCR per cohort (**Figure 2E**). The highest proportion of Mtb-specific TCR β clonotypes were found in TB diagnosis (7.5%), with a progressive decrease as individuals underwent anti-TB therapy (4.6%, 2.1%, and 1.6% at mid-treatment, end-treatment and 1-year follow up samples, respectively, **Figure 2E**). For unspecific TCR β clonotypes, the highest proportion was found in diagnosis and mid-treatment TB samples (9.2% and 9.9% respectively, **Figure 2E**), which slightly decreased in TB end-treatment and 1-year follow-up samples (6.6% respectively, **Figure 2E**). The IGRA+ cohort showed similar frequencies of Mtb-specific and unspecific TCR β clonotypes to the TB end-treatment and 1-year follow-up samples (2.3% and 5.6% respectively, **Figure 2E**).

We calculated the proportion of Mtb-specific CD4 T cells found *ex vivo* as a fraction of total number of cells sequenced in each sample (**Figure 2F**). We found the highest proportion of Mtb-specific T cells at diagnosis (2%), with a progressive decrease with treatment duration (1.3%, 0.6% and 0.4% for mid-treatment, end-treatment and 1-year post diagnosis samples, respectively, **Figure 2F**). In contrast, the proportion of unspecific CD4 T cells were lowest at diagnosis (4.6%) but highest in TB at mid treatment (7.6%), followed by end of treatment (6.5%) and one-year post diagnosis (5%) (**Figure 2F**). The IGRA+ cohort showed similar proportion of Mtb-specific CD4 T cells to the TB end-treatment cohort (0.7%), and similar frequencies of unspecific CD4 T cells to the one-year post-diagnosis and diagnosis samples (4.8%) (**Figure 2F**). Thus, by combining PDI-TCR with *ex vivo* single-cell sequencing, we successfully obtained the single-cell transcriptome of hundreds of Mtb-specific and unspecific CD4 T cells in both TB and IGRA+ cohorts.

### Mtb-specific T cells are heterogeneous and distinct between TB and IGRA+ cohorts

To characterize the transcriptome of Mtb-specific T cells, we analyzed their cell cluster composition. At diagnosis of TB, Mtb-specific T cells were predominantly found in cluster 6 (Th1*, 12% median), followed by cluster 5 (Th1, 15% median) (**Figure 3A**). In IGRA+ individuals, Mtb-specific T cells were found mostly in cluster 5 (Th1, 9 % median) or cluster 3 (memory CD127+, 9% median), but not in cluster 6 (**Figure 3B**). This indicates that Mtb-specific T cells in TB patients showed two distinct phenotypes: Th1* with high HLA-DR expression and Th1 with no HLA-DR expression. Conversely, in IGRA+ healthy individuals, Mtb-specific T cells were all HLA-DR-with either a Th1 or a memory CD127+ phenotype. Thus, distinct phenotypes of Mtb-specific T cells are linked to different disease states.

**Figure 3:**
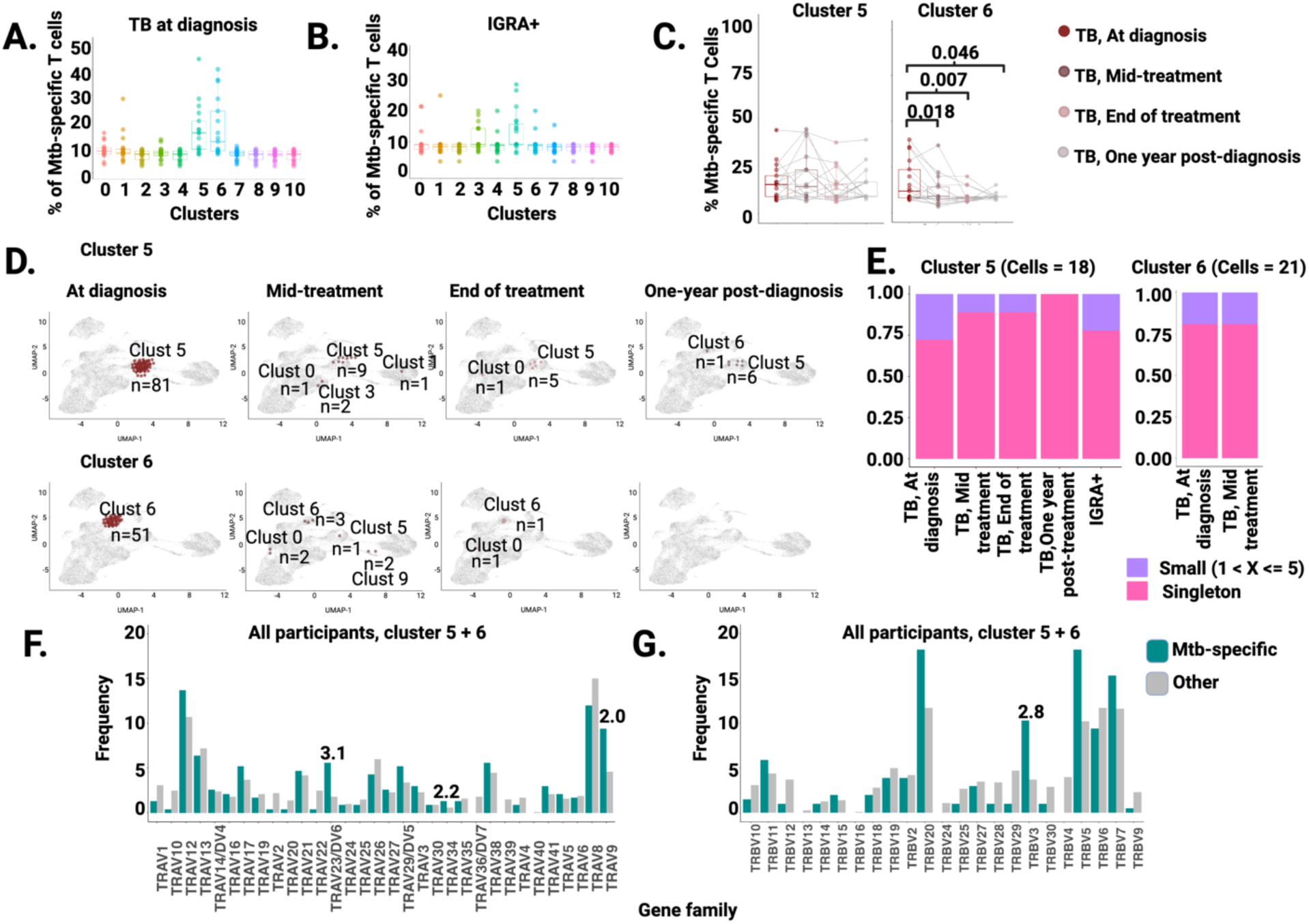
Mtb-Specific TCR Repertoire Analysis. A) Proportion of Mtb-specific cells in CD4 T cell clusters at diagnosis. This figure shows the proportion of Mtb-specific cells (y-axis) in each cluster (x-axis) at diagnosis in TB patients (n = 17). Most Mtb-specific cells were found in cluster 5 (Th1) and cluster 6 (Th1*). B) Proportion of Mtb-specific cells in CD4 T cell clusters in IGRA+ individuals. This figure shows the proportion of Mtb-specific cells (y-axis) in each cluster (x-axis) in IGRA+ individuals (n = 20). Most Mtb-specific cells were found in cluster 3 (memory CD127+) and cluster 5 (Th1). C) Proportion of Mtb-specific cells in clusters 5 (Th1) and 6 (Th1) at each visit in TB patients. This figure shows the proportion of Mtb-specific cells in cluster 5 (Th1) and cluster 6 (Th1*) at each visit in TB patients (n = 17). Dark red: TB at diagnosis; Dark pink: TB mid-treatment; Light pink: TB end of treatment; Grey: One-year post diagnosis. A significant decrease in Mtb-specific cells from diagnosis at each visit was observed in cluster 6. D) Real-time single-cell trajectory analysis of Mtb-specific cells in cluster 5 (Th1) and cluster 6 (Th1*). All clones of Mtb-specific cells found in the respective clusters at diagnosis were tracked over time to study their kinetics. At one-year post diagnosis a few cells still remained in cluster 5 but no cells were present in cluster 6. E) This figure shows the proportion of Mtb-specific clones in cluster 5 and cluster 6, down-sampled to represent an equal proportion of cells (y-axis) per cohort and visit (x-axis). Cluster 5 had 18 cells at each cohort and visit, while cluster 6 had 21 cells at diagnosis and mid-treatment. The Mtb-specific repertoire was largely composed of singletons followed by small clones (1 < x <= 5 cells). The small clones in cluster 5 were observed to decrease with treatment progression in TB patients. Pink: Singletons; Purple: small clones. F) TRA gene use frequencies in Mtb-specific vs. other TCR clonotypes in cluster 5 and 6. This figure shows the TRA gene use frequencies (y-axis) in Mtb-specific versus other TCR clonotypes in cluster 5 and cluster 6. TRAV23/DV6, TRAV34, and TRAV9 were preferentially used. Dark cyan bars: Mtb-specific TCR clonotypes in clusters 5 and 6; Grey bars: other TCR clonotypes in clusters 5 and 6. G) TRB gene use frequencies in Mtb-specific vs. other TCR clonotypes in cluster 5 and 6. This figure shows the TRB gene use frequencies (y-axis) in Mtb-specific versus other TCR clonotypes in cluster 5 and cluster 6. Only TRBV3 was preferentially used. Dark cyan bars: Mtb-specific TCR clonotypes in clusters 5 and 6; Grey bars: other TCR clonotypes in clusters 5 and 6.

### Mtb-specific T cell transcriptomes are stable but follow distinct kinetics upon anti-TB therapy

Next, we compared the effect of anti-TB therapy on the frequency and transcriptome of Mtb-specific T cells. We focused on clusters 5 and 6, which were the two clusters with the highest proportion of Mtb-specific T cells in TB diagnosis samples (**Figure 3A**). Cluster 5 showed no significant changes in the frequency of Mtb-specific T cells between diagnosis and mid-treatment, end-treatment or one-year follow up visits (**Figure 3C**, left panel). In contrast, cluster 6 exhibited a significant reduction in the proportion of Mtb-specific T cells in all samples collected after initiation of anti-TB therapy, including the mid-treatment sample, compared to the initial diagnosis sample (**Figure 3C**, right panel). Thus, distinct Mtb-specific T cell phenotypes hold distinct kinetics during anti-TB therapy.

To determine whether the transcriptome of a given clonotype was stable through treatment, we compared the transcriptome of cells expressing Mtb-specific TCRs in either cluster 5 or cluster 6 at diagnosis in the follow up visits after initiation of anti-TB therapy. While we observed a reduction in the frequency of Mtb-specific T cells initially found at diagnosis as patients underwent anti-TB therapy in both clusters, it was more pronounced in cluster 6, with no Mtb-specific T cells found at one-year post-diagnosis (**Figure 3D**). In addition, we did not observe a switch in clonotypes from one phenotype to another: for instance, Mtb-specific cells found in cluster 5 at diagnosis were predominantly also found in cluster 5 throughout the follow-up visits (**Figure 3D**). Thus, the *ex vivo* transcriptome of cells expressing a given antigen-specific TCR appears to be stable during TB therapy with no switch between effector or memory phenotypes. These data also confirm that distinct Mtb-specific T cell subsets have distinct kinetics throughout anti-TB therapy: HLA-DR+ Th1* Mtb-specific T cells show a drastic and rapid reduction with no cells left at the end of therapy, while a smaller population of HLA-DR-Th1 Mtb-specific T cells are maintained even up to six months after completion of anti-TB therapy (i.e., one-year follow-up visit).

### Mtb-specific T cells are highly diverse and are not clonally expanded

In terms of clonality, in both clusters 5 (HLA-DR-Th1) and 6 (HLA-DR+ Th1*), the Mtb-specific *ex vivo* TCR repertoire was composed of singletons or small clones and had similar composition between TB and IGRA+ samples (**Figure 3E**). There was a progressive decrease in small clones in TB patients between diagnosis and post-treatment samples in cluster 5 but not in cluster 6 (**Figure 3E**). We combined all cells expressing Mtb-specific TCRs in cluster 5 and 6 in all participants and across time points for TCR α and β chain gene usage analysis. Mtb-specific T cells had a diverse TRAV and TRBV gene usage, with a slight over-representation (i.e., 2-to 3-fold enrichment) for TRAV23/DV6, TRAV34, TRAV9 (**Figure 3F**) and TRBV3 (**Figure 3G**) compared to the other cells. Thus, both HLA-DR+ Th1* and HLA-DR-Th1 Mtb-specific T cells are highly diverse, with little clonal expansion observed even at diagnosis of TB.

### Experimental validation of Mtb-specific TCRs identified by PDI-TCR

To test if the inferred TCRs specificities in our study were accurate, we generated a Mtb-specific TCR (see methods) in a triple reporter Jurkat cell line to confirm its Mtb specificity. TCR CAYRSASDRTSGSRLTF_CSARRDSYTEAFF was selected because it had the second highest number of cells when mapped back at the single-cell level and belonged to a donor from the pilot cohort. We tested three concentrations of the Mtb peptide pool (MTB300) as well as a negative megapool (NMP, see methods). Out of the three reporters, NFAT-eGFP, NF-κB-CFP and AP-1-mCherry, only the GFP association generated a positive response (**Figure S2F**). We found a baseline response in un-pulsed and NMP samples regardless of the concentration used (5.86-17.8%). The lowest concentration of MTB300 at 0.1 µM had positive responses ranging between 11.8% - 18.1% in TCR+ cells, which was still higher compared to un-pulsed and NMP cultured cells. The next highest positive responses were observed in MTB300 at 1 µM, ranging between 20.7% - 31.0% across the three replicates. The highest responses were in MTB300 at 10 µM, ranging between 25.4% - 48.2%. This showed activation and a dose specific response in TCR+ cells only in the presence of the Mtb-peptide pool. This confirmed that the selected Mtb-specific TCR β clonotype identified by PDI-TCR with its matched TCR α chain in the *ex vivo* single cell dataset was truly specific for Mtb.

### Unspecific abundant T cells have distinct phenotype from Mtb-specific T cells

Finally, we analyzed the frequency, transcriptome and TCR repertoire of unspecific T cells using the same analysis workflow as for Mtb-specific T cells. In contrast to Mtb-specific T cells that showed heterogeneity within and across cohorts, unspecific T cells were almost exclusively found in cluster 1 (cytotoxic T cells) at diagnosis of TB (**Figure 4A**) and in IGRA+ cohorts (**Figure 4B**). During anti-TB therapy, the frequency of unspecific T cells remained constant throughout treatment (**Figure 4C**), and unspecific TCR clonotypes found in cluster 1 at diagnosis maintained a high number of cells and the same transcriptome throughout the visits (**Figure 4D**). TCR clonality analysis showed that unspecific cells were mostly large or medium expanded clones (**Figure 4E**). We combined cells expressing unspecific TCRs in cluster 1 from all participants and across timepoints for TCR α and β chain gene usage analysis. Unspecific T cells showed a diverse TRAV and TRBV gene usage and, in contrast to Mtb-specific T cells, included many genes showing over 2-fold enrichment compared to other cells in cluster 1 (**Figure 4F** and **4G**). In particular, unspecific T cells had a strong over-representation for TRAV5 and TRAV9 (17-fold and 13-fold enrichment respectively, **Figure 4F**) and TRBV18 (22-fold enrichment, **Figure 4G**). In summary, unspecific T cells were clearly distinct from Mtb-specific T cells. Unspecific T cells had a cytotoxic and highly clonally expanded phenotype, a TCR repertoire enriched for several TRAV and TRBV genes, and their frequency and transcriptome were unaffected by anti-TB therapy.

**Figure 4:**
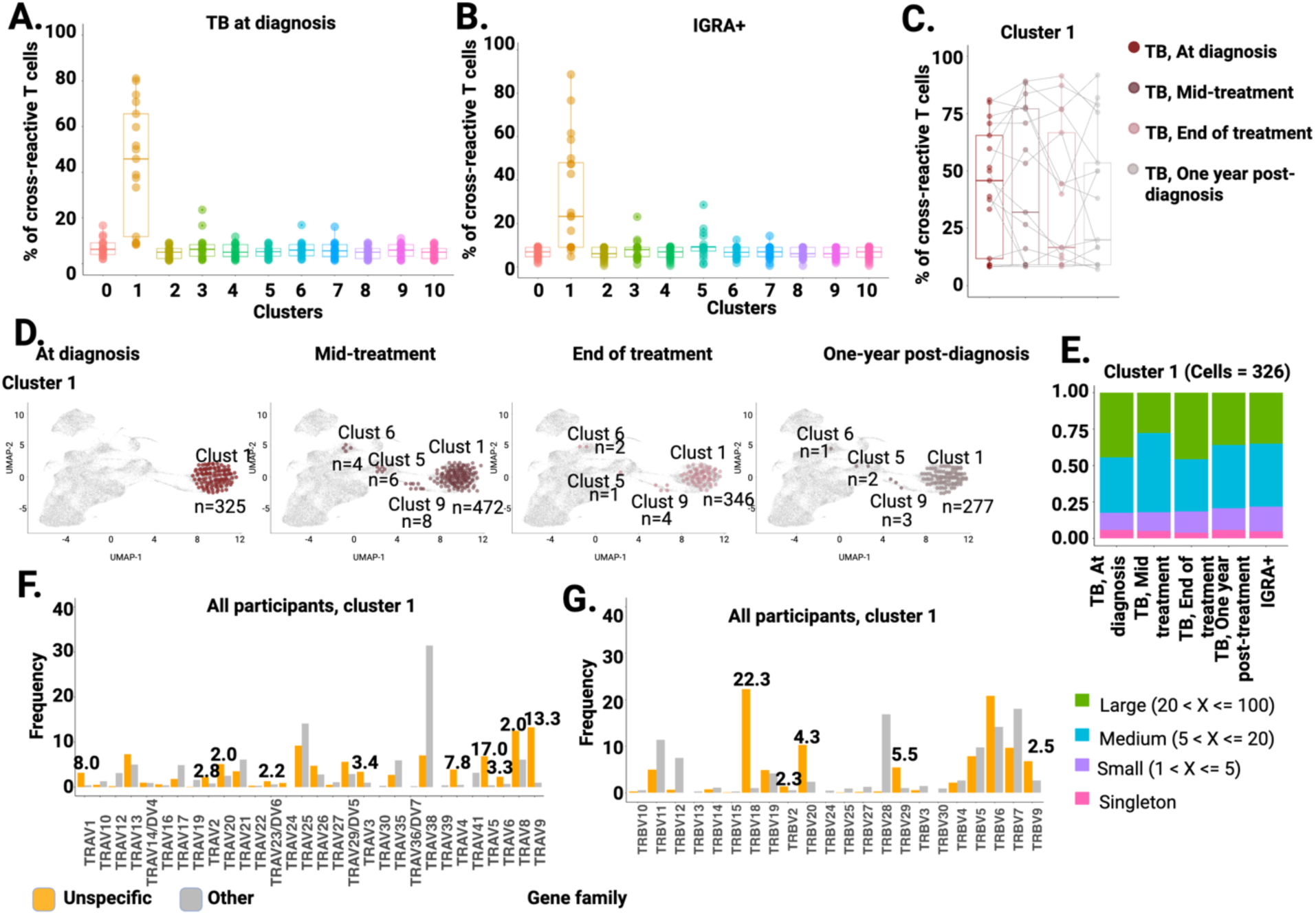
Unspecific TCR Repertoire Analysis. A) Proportion of unspecific cells in CD4 T cell clusters at diagnosis. This figure shows the proportion of unspecific cells (y-axis) in each cluster (x-axis) at diagnosis in TB patients (n = 17). Unspecific cells were mainly found in cluster 1 (cytotoxic). B) Proportion of unspecific cells in CD4 T cell clusters in IGRA+ individuals. This figure shows the proportion of unspecific cells (y-axis) in each cluster (x-axis) in IGRA+ individuals (n = 20). Unspecific cells were mainly found in cluster 1 (cytotoxic). C) Proportion of unspecific cells in clusters 1 (cytotoxic) at each visit in TB patients. This figure shows the proportion of unspecific cells in cluster 1 (cytotoxic) at each visit in TB patients (n = 17). Dark red: TB at diagnosis; Dark pink: TB mid-treatment; Light pink: TB end of treatment; Grey: One-year post diagnosis. There was no change in unspecific cell proportions with treatment. D) Real-time single-cell trajectory analysis of unspecific cells *in* cluster 1. All clones of unspecific cells found in cluster 1 at diagnosis were tracked over time to study their kinetics. Unspecific cells were remarkably stable both in their transcriptome and cell numbers over time. E) This figure shows the proportion of unspecific clones in cluster 1, down sampled to represent an equal proportion of cells (y-axis) per cohort and visit (x-axis). Cluster 1 had 326 cells at each cohort and visit. The unspecific repertoire was largely composed of large (20 < x <= 100) and medium clones (5 < x <= 20 cells). Green: Large clones; Blue: Medium clones; Purple: Small clones; Pink: Singletons. F) TRA gene use frequencies in unspecific vs. other TCR clonotypes in cluster 1. This figure shows the TRA gene use frequencies (y-axis) in unspecific versus other TCR clonotypes in cluster 1. TRAV5 and TRAV9 had high use. Yellow bars: Unspecific TCR clonotypes in clusters 1; Grey bars: other TCR clonotypes in cluster 1. G) TRB gene use frequencies in unspecific vs. other TCR clonotypes in cluster 1. This figure shows the TRB gene use frequencies (y-axis) in unspecific versus other TCR clonotypes in cluster 1. TRBV18 was highly used. Yellow bars: Unspecific TCR clonotypes in cluster 1; Grey bars: other TCR clonotypes in cluster 1.

## Discussion

In this study, we developed the **<u>P</u>**eptide-**<u>D</u>**riven **<u>I</u>**dentification of **<u>TCR</u>**s (PDI-TCR) assay, which utilizes *in vitro* stimulation with antigen-specific peptide pools, bulk TCR sequencing and a statistical analysis workflow to identify antigen-specific TCR β clonotypes. Its primary distinction from previous methods lies in the incorporation of a control culture stimulated with a peptide pool for a commonly encountered antigen (distinct from the one to be studied) and a stringent pipeline to confidently identify “true” antigen-specific and unspecific TCR β clonotypes. There are several advantages to using PDI-TCR over other methods for identifying antigen-specific TCRs: first there is no need for previous knowledge of HLA/peptide combinations as with tetramers. Second, it does not rely on phenotyping and isolating antigen-specific T cells using AIM markers and cell sorting. Third, it is tailored to work with samples with small blood volumes (i.e., 5 to 10 mL of blood) which can often be a limiting factor. Finally, by using a peptide pool targeted to a common pathogen in parallel to the peptide pool of interest, PDI-TCR can identify both antigen-specific and unspecific TCRs.

By applying PDI-TCR on a pilot cohort of four TB patients with Mtb- and BP-specific *in vitro* culture replicates and *ex vivo* samples, we showed that the ability of PDI-TCR to identify a given antigen-specific TCR clonotype repeatedly was directly dependent on its *ex vivo* abundance. This is because more prevalent clonotypes are more likely to be present in the starting pool of PBMC used in each of the two independent replicates. In addition, among the repeated TCR β clonotypes, we found comparable proportions of Mtb-specific and unspecific TCR β clonotypes but no BP-specific TCR β clonotypes. Given that the assay was conducted on samples from patients with TB but not *B. pertussis* infection, we expected a higher frequency of circulating Mtb-specific T cells over BP-specific T cells, thus more repeated Mtb-specific TCR clonotypes across PDI-TCR replicates. Together, this indicates that PDI-TCR may be most suited for studying active/recent infections or vaccine booster studies, where a relatively high abundance of antigen-specific memory T cells is expected to be found in PBMC.

We applied PDI-TCR to a larger cohort of 19 TB patients and 19 IGRA+ individuals and identified hundreds of Mtb-specific and unspecific TCR clonotypes in each cohort. We also performed single-cell RNA and TCR sequencing on *ex vivo* sorted CD4 T cells in the same samples and analyzed the transcriptome of Mtb-specific and unspecific T cells directly *ex vivo*. This is significant, as the only other technique in common use to isolate *ex vivo* antigen specific cells require MHC:epitope tetramers (or the like) to sort cells, and generating such tetramer reagents for a broad population and an epitope set is not feasible. Other techniques that bypass tetramers to identify antigen-specific TCR rely on *in vitro* stimulation, which alters the T cell transcriptome compared to direct *ex vivo* sampling. Only a small fraction of Mtb-specific and unspecific TCR β clonotypes identified with PDI-TCR were also found in the *ex vivo* single-cell dataset (1.6-7.5% of Mtb and 5.6-9.9% of unspecific). This is likely due to too few cells sequenced *ex vivo*, and we hypothesize that if we had sequenced more cells, we could have obtained a larger repertoire and thus mapped more Mtb-unspecific T cells *ex vivo*.

We found that Mtb-specific T cells had a highly diverse TCR repertoire, and transcriptomic heterogeneity within and across cohorts. In TB, Mtb-specific T cells showed two distinct transcriptomes and kinetics throughout treatment. Mtb-specific T cells with an HLA-DR+ Th1* phenotype showed significant decrease during treatment, with no detectable cells at the end of anti-TB therapy. In contrast, Mtb-specific T cells with an HLA-DR-Th1 phenotype retained a smaller but steady population post treatment. In IGRA+, Mtb-specific T cells displayed HLA-DR-Th1 and memory CD127+ phenotypes, but no HLA-DR+ Th1* phenotypes. We and others have previously shown than HLA-DR marks Mtb-specific effector T cells in TB patients at diagnosis but not IGRA+ individuals^40,46,47^. Our new results confirm this observation, and further advance that HLA-DR+ Th1* Mtb-specific T cells likely represent short-lived effector T cells, while Mtb-specific T cells with an HLA-DR-Th1 phenotype represent a subset of longer-lived memory T cells.

In terms of Th subsets, previous work from our group and others have shown that Mtb-specificity resides predominantly in Th1* cells in both IGRA+ individuals^31,48,49^ and TB patients^50^. In contrast, in our dataset we found that Mtb-specific T cells with a Th1* phenotype were exclusively present in TB patients. This failure to identifying Th1* cells in IGRA+ individuals where they were first defined is likely due to the difference in defining cell phenotypes based on the expression of chemokine receptors on the cell surface by flow cytometry (as previous studies did) vs. the *ex vivo* transcriptome of these cells. Thus, the CXCR3+CCR6+CCR4-definition of Th1* by flow cytometry may encompass different T cell subsets, some resembling Th1*, and other closer to Th1 at the transcriptomic level. It would be informative to repeat the *ex vivo* single-cell experiment using a panel of CITE-Seq antibodies including chemokine receptor markers typically used in flow cytometry to identify Th1 and Th1*.

Surprisingly, we did not find that TCR clonotypes altered their gene expression during anti-TB therapy. Mtb-specific TCR clonotypes present in the effector-like Th1* HLA-DR+ cluster in TB at diagnosis did entirely disappear from the global CD4 T cell pool in the subsequent visits and were not found in other clusters. In addition, Mtb-specific T cells were small clones or singletons, and we saw no changes in TCR clonality across cohorts and visits. This was a surprising find, as we were expecting most effector T cells to be clonally expanded in TB at diagnosis, and some effector cells to revert to memory populations as individuals underwent treatment. It is possible that the effector to memory transition was indeed occurring but because the remaining memory clones were so few and rare, our dataset did not have adequate sequencing depth to detect them. In terms of clonality, similarly, the pool of CD4 T cell sequenced *ex vivo* may have been too small to identify TCR clonotypes that had small expansions.

In addition to Mtb-specific TCR β clonotypes, the PDI-TCR workflow allowed us to identify unspecific TCR β clonotypes that were present at high frequencies in both Mtb and BP cultures. Although we named these clonotypes “unspecific”, our statistical definition did not take in consideration the *ex vivo* abundance of each clonotypes, and thus included TCR β clonotypes that were both truly expanded between *ex vivo* and *in vitro* stimulation (i.e., cross-reactive), versus those with constitutive high abundance. This highlights that without a corresponding *ex vivo* sample, PDI-TCR is not able to differentiate between these two subcategories of unspecific T cells.

However, we found that even when using our less granular definition, unspecific T cells were remarkably homogenous across cohorts, and clearly distinct from Mtb-specific T cells.

Unspecific T cells were found to be cytotoxic, highly expanded in both TB and IGRA+ individuals, and had enrichment for specific TRA and TRB genes, thus indicating reduced repertoire diversity compared to Mtb-specific T cells. They were also minimally affected by anti-TB therapy. We found no association in the literature to the highly used TRA and TRB genes by Mtb- or unspecific TCRs. Interestingly TRAV9 was preferentially used by both Mtb- and unspecific TCR clonotypes suggesting that it could be a frequently used gene with no links to specificity or cell type.

Taken together, our results show that unspecific T cells present several features of unconventional CD4 T cells, that may not rely on peptide/MHC/TCR interactions for activation^51,52,53,54,55^. Importantly, unspecific T cells represented a non-negligible fraction of CD4 T cells retrieved after stimulation in PDI-TCR as well as *ex vivo*, with similar or higher abundance than Mtb-specific T cells. This underscores the importance of using a non-related antigen stimulation as a control for PDI-TCR, as well as in other antigen-specific TCR assays that relies on *in vitro* stimulation (e.g., AIM assay) to prevent unspecific T cell from being mislabeled as antigen-specific.

In summary, we developed PDI-TCR as a novel method to confidently identify antigen-specific TCR β clonotypes to a given pathogen, as well unspecific TCR β clonotypes between non-related pathogens. When combined with single-cell TCR and RNA sequencing, PDI-TCR provided a unique opportunity to characterize the *ex vivo* transcriptome of antigen-specific T cells, and importantly to follow a given TCR clonotype over time in longitudinal samples. Using this method, we found previously uncharacterized heterogeneity within Mtb-specific T cells in TB, including a unique phenotype exclusively found prior to anti-TB therapy and resembling short-lived effector T cells. Additionally, our study uncovered the presence of unspecific T cells that are highly homogenous across cohorts with features resembling unconventional T cells, including high clonal expansion, cytotoxic phenotype, and restricted TCR repertoire.

Collectively, our findings highlight the potential of PDI-TCR, in conjunction with single-cell sequencing, as a powerful tool for studying antigen-specific T cells.

## Data availability

The processed data from bulk and single-cell sequencing has been submitted to NCBI (National Center for Biotechnology Information; https://www.ncbi.nlm.nih.gov/) and can be found under the submission ID: GSE293807.

## Study Funding

This work was supported by the National Institutes of Health grant U19 AI118626 (to B.P.), National Institutes of Health contract 75N93024C00057 (to B.P. and C.S.L.A) and National Science Foundation Fellowship grant DGE-2038238 (to L.Y.C.). The funders had no role in study design, data collection, analysis, decision to publish, or manuscript preparation.

## Technical Support

We sincerely appreciate the support of the Vijayanand Lab in sequencing samples, as well as the assistance provided by the Flow Cytometry and Bioinformatics Cores at the La Jolla Institute for Immunology in sample acquisition and data analysis.

## Competing Interests

The authors have declared that no competing interests exist.

## Author Contributions

J.B. and B.P. participated in the design and direction of the study. R.T., J.B., L.Y.C., C.S.L.A., and B.P. performed and analyzed the experiments. R.T., R.T., J.B. and B.P. performed bioinformatics analysis. A.C., K.F., J.G., K.K. assisted in bioinformatics analysis. B.P., C.S.L.A., and A.S. provided peptide pools and reagents for experiments. R.T., J.P., H.G., D.D.S., W.G., T.J.S. and A.D.DS, recruited participants, performed clinical evaluations, and isolated PBMCs. R.T., J.B., L.Y.C., C.S.L.A. and B.P. wrote the manuscript. All authors read, edited, and approved the manuscript before submission.

## Notes

https://www.ncbi.nlm.nih.gov/

### Competing Interest Statement

The authors have declared no competing interest.

