## Supplementary material for "Peptide Driven Identification of TCRs (PDI-TCR) reveals dynamics and phenotypes of CD4 T cells in tuberculosis": Sup Figures and Tables

### Supplementary Figures

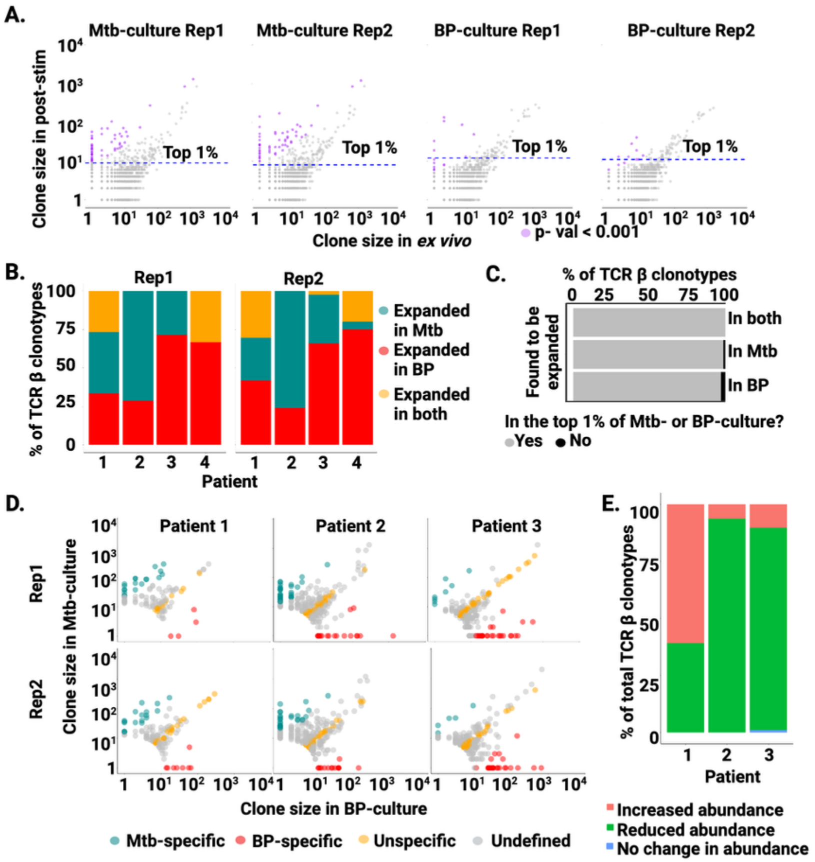


**Figure S1: Analysis of TCR β Clonotype Abundance in Mtb- vs BP- Cultures.**

A) Representative plots from one patient for both replicates for all TCR β clonotypes of Mtb- or BP-cultures versus *ex vivo*. Each dot represents a TCR β clonotype plotted by their abundance in Mtb-culture or BP-culture (y-axis) versus *ex vivo* (x-axis). Purple: TCR β

Clonotypes with a p-value <0.001; TCR β clonotypes above the dotted line are in the top 1% of abundance in Mtb- or BP- culture. B) Proportion of TCR β clonotypes that were found to be significantly expanded (p-value < 0.001) in Mtb- (dark cyan), BP- (red) or both (yellow) compared to *ex vivo* in all four TB patients tested in the pilot experiment. C) Proportion of significantly expanded TCR β clonotypes found in the top 1% of abundance in each patient (n = 4) in Mtb-cultures, BP-cultures, or both. Significantly expanded TCR β clonotypes had a p-value < 0.001 in post-stimulation versus *ex vivo*. Grey: TCR β clonotypes in the top 1%; Black: TCR β clonotypes not in the top 1%. D) Representative plots for the top 1% of the most abundant TCR β clonotypes subjected to Fisher's exact test in both replicates of Mtb- versus BP- in all three patients that showed TCR β clonotypes expanded in Mtb-cultures. TCR β clonotypes with a p-value <0.001 and a FC cutoff >10 in Mtb- vs BP- cultures were selected. Each dot represents a TCR β clonotype. Mtb-specific TCR β clonotypes are indicated in dark cyan, BP-specific TCR β clonotypes in red, and unspecific TCR β clonotypes in yellow. E) Proportion of unspecific TCR β clonotypes that increased, decreased, or remained unchanged in abundance after a 14-day expansion in culture, compared to their *ex vivo* levels. The data is presented as the percentage of total unspecific TCR β clonotypes identified in all three patients that showed TCR β clonotypes expanded in Mtb-cultures. Pink: TCR β clonotypes that increased in abundance compared to *ex vivo*; Green: TCR β clonotypes that decreased in abundance compared to *ex vivo*; Blue: TCR β clonotypes that did not change in abundance from *ex vivo*.

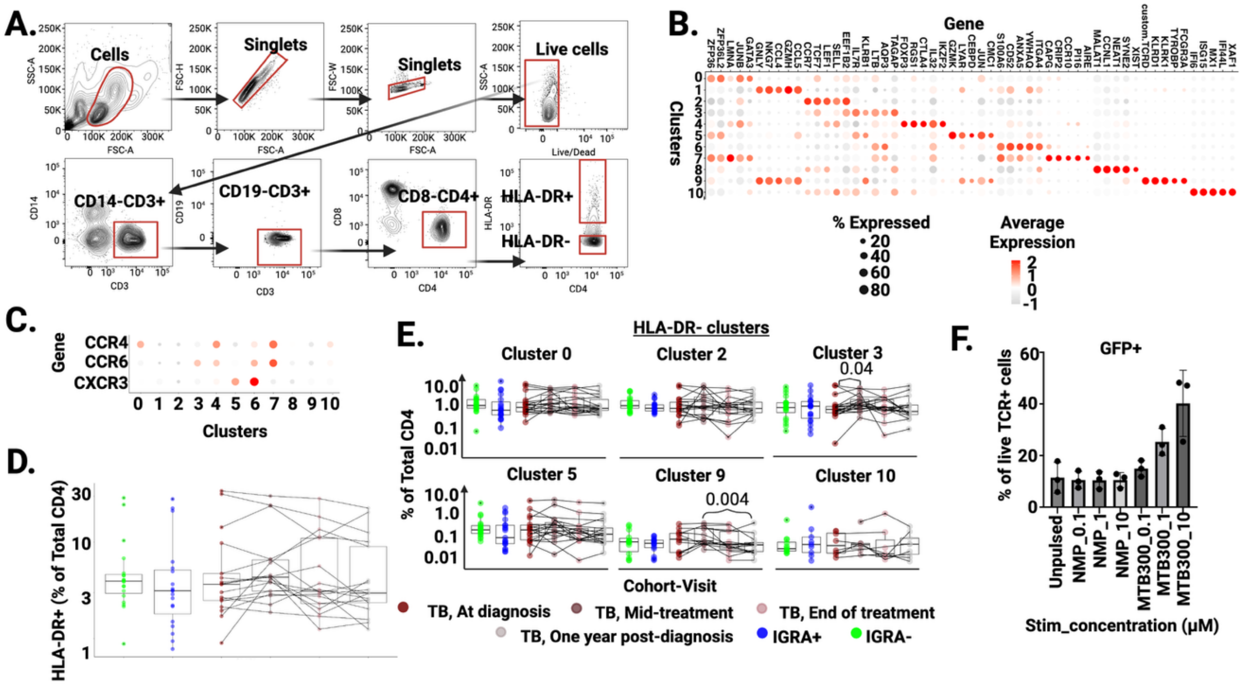


**Figure S2: HLA-DR Marks Mtb-Specific Activation.**

A) Gating strategy to sort *ex vivo* HLA-DR+ and HLA-DR- CD4 T cells in human PBMCs by flow cytometry in TB, IGRA+ and IGRA- cohorts. B) Dot plot of the top 5 markers in each cluster arranged by $log_2$FC. Size of each dot is the fraction of cells expressing the feature. The color scale represents the expression level. C) Dot plot of CXCR3, CCR6 and CCR4 gene expression. The size of each dot is the fraction of cells expressing the feature, the color scale represents the expression level. D) Percent of total HLA-DR+ CD4 T cells in each sample per donor as determined by flow cytometry (n = 57). E) Adjusted cell counts in HLA-DR- CD4 T cell clusters (clusters 0, 2, 3, 5, 9 and 10). The number of cells in each cluster was adjusted to reflect the true frequency of HLA-DR- cells in total CD4 T cells (y-axis) within each sample per donor (n=57) based on the flow cytometry data. F) NFAT-eGFP Reporter expression in Jurkat cells expressing a Mtb-specific TCR, performed in triplicate. The cells were co-cultured with a lymphoblastoid cell line (LCL) that was either un-pulsed or pulsed with 0.1, 1, or 10 μM of a Negative Mega Pool (NMP) or 0.1, 1, or 10 μM of Mtb peptides (MTB300).

### Supplementary Tables

**Table S1: TotalSeq antibodies**

| Barcode | Specificity | Clone | Manufacturer | Volume in ul added to 100 ul of staining volume | Category |
|---|---|---|---|---|---|
| 159 | HLA-DR | L243 | BioLegend | 2 | TotalSeqTM-C |
| 45 | CD4 | SK3 | BioLegend | 2 | TotalSeqTM-C |
| 48 | CD45 | 2D1 | BioLegend | 2 | TotalSeqTM-C |
| 391 | CD45 | HI30 | BioLegend | 2 | TotalSeqTM-C |
| 251 | CD298 and β2 microglobulin | LNH-94; 2M2 | BioLegend | 2 | TotalSeqTM-C |
| 252 | CD298 and β2 microglobulin | LNH-94; 2M2 | BioLegend | 2 | TotalSeqTM-C |
| 253 | CD298 and β2 microglobulin | LNH-94; 2M2 | BioLegend | 2 | TotalSeqTM-C |
| 254 | CD298 and β2 microglobulin | LNH-94; 2M2 | BioLegend | 2 | TotalSeqTM-C |
| 255 | CD298 and β2 microglobulin | LNH-94; 2M2 | BioLegend | 2 | TotalSeqTM-C |
| 256 | CD298 and β2 microglobulin | LNH-94; 2M2 | BioLegend | 2 | TotalSeqTM-C |
| 257 | CD298 and β2 microglobulin | LNH-94; 2M2 | BioLegend | 2 | TotalSeqTM-C |
| 258 | CD298 and β2 microglobulin | LNH-94; 2M2 | BioLegend | 2 | TotalSeqTM-C |
| 259 | CD298 and β2 microglobulin | LNH-94; 2M2 | BioLegend | 2 | TotalSeqTM-C |

| | | | | | |
|---|---|---|---|---|---|
| 260 | CD298 and β2 microglobulin | LNH-94; 2M2 | BioLegend | 2 | TotalSeqTM-C |

**Table S2: Surface antibody panel for flow cytometry**

| Antibody | Fluorochrome | Clone | Volume in ul added to 100 ul of staining volume | Manufacturer |
|---|---|---|---|---|
| CD45 | BUV395 | HI30 | 5 | BD Biosciences |
| CD8 | BUV496 | RPA-T8 | 5 | BD Biosciences |
| CD20 | BUV563 | 2H7 | 5 | BD Biosciences |
| CD16 | BUV737 | 3G8 | 5 | BD Biosciences |
| CD3 | BUV805 | UCHT1 | 5 | BD Biosciences |
| CD14 | BV480 | M5E2 | 5 | BD Biosciences |
| CD45RA | BV570 | HI100 | 5 | BioLegend |
| CD19 | BV605 | HIB19 | 5 | BioLegend |
| IgD | BV750 | IA6-2 | 5 | BD Biosciences |
| CD11c | BV785 | 3.9 | 5 | BioLegend |
| CCR7 | BB515 | 2-L1-A | 5 | BD Biosciences |
| CD123 | PerCP-Cy5.5 | 6H6 | 5 | BioLegend |
| CD38 | PE-Dazzle594 | HIT2 | 5 | BioLegend |
| HLA-DR | PE-Cy7 | LN3 | 2 | BioLegend |
| CD56 | APC | 5.1H11 | 5 | BioLegend |
| CD4 | APC-eF780 | RPA-T4 | 5 | eBioscience |
| CD71 | PE-Cy5 | M-A712 | 5 | BD Biosciences |
| TCRgd | BV421 | 11F2 | 4 | BD Biosciences |
| CD1c | BV650 | L161 | 5 | BioLegend |

**Table S3: Total β sequence reads**

| Patient | Condition | Replicate | #Cells | #TCR β clonotypes | #Singletons | % of Singletons |
|---|---|---|---|---|---|---|
| Patient_1 | *Ex vivo* | N/A | 42,990 | 27,287 | 21,934 | 80 |
| | Mtb-specific stimulation | Rep1 | 13,225 | 6,935 | 5,562 | 80 |
| | Mtb-specific stimulation | Rep2 | 12,930 | 7,225 | 5,817 | 81 |
| | BP-specific stimulation | Rep1 | 8,863 | 5,855 | 4,896 | 84 |
| | BP-specific stimulation | Rep2 | 10,709 | 6,886 | 5,731 | 83 |
| Patient_2 | *Ex vivo* | N/A | 85,866 | 56,882 | 49,311 | 87 |
| | Mtb-specific stimulation | Rep1 | 23,884 | 13,400 | 11,587 | 86 |
| | Mtb-specific stimulation | Rep2 | 25,680 | 14,457 | 12,482 | 86 |
| | BP-specific stimulation | Rep1 | 24,358 | 16,176 | 14,324 | 89 |
| | BP-specific stimulation | Rep2 | 21,711 | 15,451 | 13,487 | 87 |
| Patient_3 | *Ex vivo* | N/A | 104,496 | 59,912 | 51,250 | 86 |
| | Mtb-specific stimulation | Rep1 | 18,235 | 11,506 | 9,978 | 87 |
| | Mtb-specific stimulation | Rep2 | 20,796 | 12,324 | 10,561 | 86 |
| | BP-specific stimulation | Rep1 | 22,352 | 13,581 | 11,698 | 86 |
| | BP-specific stimulation | Rep2 | 21,399 | 13,295 | 11,411 | 86 |
| Patient_4 | *Ex vivo* | N/A | 32,698 | 27,231 | 24,603 | 90 |
| | Mtb-specific stimulation | Rep1 | 7,394 | 5,659 | 5,176 | 91 |
| | Mtb-specific stimulation | Rep2 | 11,238 | 8,674 | 7,935 | 91 |
| | BP-specific stimulation | Rep1 | 7,653 | 6,093 | 5,613 | 92 |
| | BP-specific stimulation | Rep2 | 10,049 | 7,645 | 7,024 | 92 |

**Table S4: Summary statistics of the Fisher's exact test**

| Sample | # Total TCR β clonotypes | # Top 1% TCR β clonotypes | p-value < 0.05 | | | p-value < 0.01 | | | p-value < 0.001 | | |
|---|---|---|---|---|---|---|---|---|---|---|---|
| | | | # Positives | # EFP | FDR % | # Positives | # EFP | FDR % | # Positives | # EFP | FDR % |
| Patient 1 Rep1 | 11550 | 115 | 69 | 5.75 | 8.33 | 55 | 1.15 | 2.09 | 38 | 0.1 | <1 |
| Patient 1 Rep2 | 12748 | 127 | 58 | 6.35 | 10.9 | 49 | 1.27 | 2.59 | 33 | 0.1 | <1 |
| Patient 2 Rep1 | 28040 | 280 | 129 | 14 | 10.9 | 104 | 2.8 | 2.69 | 76 | 0.3 | <1 |
| Patient 2 Rep2 | 28492 | 284 | 154 | 14.2 | 9.22 | 117 | 2.84 | 2.43 | 78 | 0.3 | <1 |
| Patient 3 Rep1 | 23406 | 234 | 68 | 11.7 | 17.2 | 52 | 2.34 | 4.5 | 43 | 0.2 | <1 |
| Patient 3 Rep2 | 23830 | 238 | 81 | 11.9 | 14.7 | 63 | 2.38 | 3.78 | 46 | 0.2 | <1 |
| Patient 4 Rep1 | 11290 | 112 | 3 | 5.6 | 187 | 1 | 1.12 | 112 | 0 | 0.1 | N/A |
| Patient 4 Rep2 | 15623 | 156 | 4 | 7.8 | 195 | 1 | 1.56 | 156 | 1 | 0.2 | N/A |

**Table S4: Top 50 markers in each cluster** (provided as an Excel file)
